# Efferocytosis-Induced Lactate Enables the Proliferation of Pro-Resolving Macrophages to Mediate Tissue Repair

**DOI:** 10.1101/2023.03.10.532107

**Authors:** David Ngai, Maaike Schilperoort, Ira Tabas

## Abstract

The clearance of apoptotic cells (ACs) by macrophages (efferocytosis) promotes tissue repair by preventing necrosis and inflammation and by activating pro-resolving pathways, including continual efferocytosis. A key resolution process *in vivo* is efferocytosis-induced macrophage proliferation (EIMP), in which AC-derived nucleotides trigger Myc-mediated macrophage proliferation, thereby increasing the pool of efferocytosis-competent macrophages. Here we show that EIMP requires a second input that is integrated with cellular metabolism, notably, efferocytosis-induced lactate production. While the AC-nucleotide pathway leads to induction of *Myc* mRNA, lactate signaling is required for the stabilization of Myc protein and subsequent macrophage proliferation. Lactate, via GPR132 and protein kinase A, activates AMP kinase, which increases the NAD^+^:NADH ratio. This upstream pathway then activates the NAD^+^-dependent protein deacetylase, SIRT1, which deacetylates Myc to promote its stabilization. Inhibition or silencing of any step along this pathway prevents the increase in Myc protein and proliferation in efferocytosing macrophages despite the presence of the AC-nucleotide/*Myc* mRNA pathway. To test importance *in vivo*, we transplanted mice with bone marrow cells in which the lactate biosynthetic enzyme lactate dehydrogenase A (LDHA) was knocked down. We then subjected these mice and control bone marrow-transplanted mice to dexamethasone-induced thymocyte apoptosis, a model of high-AC burden. The thymi of the LDHA-knockdown cohort showed reduced macrophage Myc protein expression and proliferation, impaired AC clearance, and increased tissue necrosis. Thus, efferocytosis-induced macrophage proliferation, which is a key process in tissue resolution, requires inputs from two independent efferocytosis-induced processes: a signaling pathway induced by AC-derived nucleotides and a cellular metabolism pathway involving lactate production. These findings illustrate how seemingly distinct pathways in efferocytosing macrophages are integrated to carry out a key process in tissue resolution.

## Introduction

The clearance of apoptotic cells (ACs), known as efferocytosis, is a critical process mediated mostly by macrophages to promote tissue repair and homeostasis^1–4^. Efferocytosis prevents pro-inflammatory secondary necrosis and promotes the secretion of pro-resolving factors, such as TGF-β and IL-10, to suppress inflammation and promote tissue resolution^1–4^. Pro-resolving factors can, in turn, promote efferocytosis, particularly the sequential uptake of multiple ACs by a single macrophage known as continual efferocytosis^5^. This process forms a physiologically important efferocytosis-resolution positive-feedback cycle^1–4^.

Macrophage functions, including those involved in efferocytosis and resolution, is profoundly affected by intracellular metabolism and metabolic by-products. For example, lactate produced by glycolytic metabolism plays an active role in pro-resolution signaling in immune cells, including macrophages^6–9^. Two recent studies have shown that efferocytosis can drive increased glycolysis to produce lactate, which stimulates macrophage secretion of pro-resolving factors and promotes continual efferocytosis^8,9^. Another form of macrophage metabolism that occurs during efferocytosis is “cargo” metabolism, whereby metabolites released during phagolysosomal degradation of ACs, such as nucleic acids^10–12^, can trigger pro-resolving pathways and continual efferocytosis in efferocytosing macrophages. A physiologically important example of this principle is a macrophage proliferation pathway activated by AC-derived nucleotides, which functions *in vivo* to promote AC clearance and tissue resolution by expanding the pool of pro-resolving macrophages^10^. This process, termed efferocytosis-induced macrophage proliferation (EIMP), involves two convergent pathways: 1) an ERK1/2 pathway activated when ACs engage the macrophage MerTK receptor; and 2) a DNA-PK-mTORC2-Akt pathway activated by AC-derived oligonucleotides after the engulfed ACs are degraded in phagolysosomes. These pathways result in increased transcription of *Myc* mRNA, leading to Myc-mediated cell cycling^10^. As Myc protein stability is an important determinant of Myc protein levels^13–19^, we wondered how this aspect of Myc regulation factored into EIMP.

Glycolytic metabolism and lactate have been shown to promote proliferation in other cell types, such as vascular smooth muscle cells and regulatory T-cells^20,21^. Moreover, lactate has been linked to the activation of the NAD^+^-dependent protein deacetylase, SIRT1, which is a protein deacetylase known to deacetylate Myc and promote its stability^14–16^. We therefore hypothesized that efferocytosis-induced lactate, which we call EIL, could promote EIMP through the stabilization of Myc protein by SIRT1-mediated deacetylation. In support of this idea, we show that lactate production during efferocytosis is necessary to increase Myc protein through protein stabilization and is required for EIMP. Myc stabilization requires SIRT1, which is downstream of a PKA-AMPK-NAD^+^ pathway, and PKA is activated by secreted lactate acting through the G protein-coupled receptor 132 (GPR132). Inhibition or genetic targeting of the rate-limiting enzyme for lactate production, lactate dehydrogenase A (LDHA), reduces Myc protein and EIMP both *in vitro* and *in vivo,* leading to impaired macrophage proliferation and AC clearance and increased tissue necrosis. Thus, distinct metabolic pathways in efferocytosing macrophages, one involving AC cargo and the other involving lactate production, are integrated to enable the expansion of pro-resolving macrophages and subsequent tissue repair.

## Methods

### Cell lines

L-929 fibroblasts and Jurkat T-lymphocytes were obtained from ATCC. Both cell lines were cultured in DMEM supplemented with 10% heat-inactivated fetal bovine serum (HI-FBS) and 100 U/mL penicillin-streptomycin (Gibco). Cells were incubated at 37°C with 5% CO_2_.

### Experimental animals

Animal protocols used were approved by Columbia University’s Institutional Animal Care and Use Committee and were cared for according to NIH guidelines for the care and use of laboratory animals. The mice were socially housed in standard cages at 22°C and with a 12-hour light-dark cycle in a barrier facility with ad libitum access to food and water. 8-10 week-old male C57BL/6J mice (JAX 000664) were used as bone marrow-transplantation (BMT) recipient mice for the dexamethasone-thymus experiment. *Ldha^fl/fl^* mice used as bone marrow donors were 8–10-week-old male mice on the C57BL/6J background (JAX 030112).

### Bone marrow transplantation

8-10 week-old C57BL/6J male mice (JAX) were irradiated with 10 Gγ with a cesium-137 γ-emitting irradiator (Gamma cell 40, MSD Nordion). Bone marrow cells harvested from *Ldha^fl/fl^* mice (JAX 030112) were incubated with 5 μM TAT-CRE Recombinase (Millipore) for 30 minutes to delete the *Ldha* gene. Bone marrow cells were then rinsed twice with PBS. *Ldha^fl/fl^* bone marrow cells not treated with TAT-Cre Recombinase were used as the control. 4 hours after irradiation, 2.5 x 10^6^ bone marrow cells were administered to recipient mice by tail vein injection. Recipient mice were allowed to recover for 4 weeks and given water containing 10 mg/mL neomycin for the first 3 weeks. These mice were then used for the dexamethasone-thymus assay.

### Dexamethasone-thymus assay

Mice transplanted with control or *Ldha*-deleted bone marrow were injected i.p. with 250 μg dexamethasone (Sigma) in PBS. Eighteen hours after injection, the thymi were harvested, fixed in 10% formalin, embedded in paraffin, and sectioned at a thickness of 5 μm. Sections were stained as described below under Immunofluorescence staining and microscopy. Efferocytosis was quantified as the ratio of Mac2^+^-associated TUNEL^+^ cells to free TUNEL^+^ cells. Macrophage proliferation was assessed by quantifying Ki67^+^ Mac2^+^ cells that were TUNEL^+^ or TUNEL^-^. Myc expression was assessed by measuring Myc MFI in Mac2^+^ TUNEL^+^ or Mac2^+^ TUNEL^-^ cells. To measure necrotic area, H&E-stained thymic sections were imaged and regions with hypocellularity and fragmented nuclei were quantified.

### Mouse bone marrow-derived macrophages (BMDMs)

Bone marrow cells were isolated from the femurs of 8-10 week old male C57BL/6J mice, *Ldha^fl/fl^* mice, or *Ldha^fl/fl^*; *LysMCre*^+/-^ mice and were cultured for 7 days in BMDM differentiation medium, which was DMEM supplemented with 10% (v/v) heat-inactivated fetal bovine serum (HI-FBS); 100 U/mL penicillin and 100 U/mL streptomycin (Corning); and 20% (v/v) L-929 cell-conditioned medium. Femurs from *Ldha^fl/fl^* mice and *Ldha^fl/fl^*; *LysMCre^+/-^* mice were generously provided by Dr. Lev Becker (University of Chicago). Cells were incubated at 37°C in a 5% CO_2_ incubator.

### Human monocyte-derived macrophages (HMDMs)

Peripheral blood mononuclear cells were isolated from the buffy coats of anonymous healthy adult donors with informed consent (New York Blood Center) using Histopaque-1077 (Sigma). Isolated cells were rinsed and cultured for 7-14 days in RPMI-1640 medium with L-glutamine (10-040; Corning) supplemented with 10% (v/v) HI-FBS (Gibco), 10 U/mL penicillin and 100 U/mL streptomycin (Corning), and 10 ng/mL GM-CSF (PeproTech).

### Generation of ACs and incubation with macrophages

Jurkat cells were resuspended in PBS and irradiated with a 254-nm UV lamp for 15 minutes to induce apoptosis. To fluorescently label ACs, irradiated ACs were pelleted by centrifugation, resuspended in Diluent C (Sigma), mixed with Diluent C containing PKH26 (red) (Sigma), and incubated at 37°C for 5 minutes. The labeling reaction was halted with an equal volume of HI-FBS. ACs were pelleted by centrifugation, resuspended in PBS, and incubated at 37°C for 2-3 hours. The ACs were collected by centrifugation, resuspended in fresh PBS, added to macrophages at a 5:1 AC:macrophage ratio, and incubated for 45 minutes at 37°C. The macrophage monolayers were then rinsed with PBS to remove unbound ACs and chased for 1, 3, or 24 hours in full medium with or without the addition of 50 μM FX11 (Sigma), 10 μM EX-527 (Sigma), 10 μM Compound C (Sigma), 10 μM H-89 (Sigma), 10 μM MG-132 (Sigma), 10 mM lactic acid (Sigma), 500 μM NMN (Sigma), 10 μM telmisartan (MedChemExpress), or 20 μM ONC-212 (MedChemExpress).

### Lactate assay

Macrophages were seeded at 200,000 cells per well in 24-well plates in BMDM differentiation medium. After AC incubation, macrophages were chased for 1 hour in 200 μL of low-serum DMEM (DMEM supplemented with 1% (v/v) HI-FBS) + 50 μM FX11. Media was collected and assayed using a lactate assay kit from Sigma-Aldrich (MAK064) according to the manufacturer’s instructions.

### siRNA transfection

Macrophages were seeded at ~60% confluency in BMDM differentiation media. Scrambled siRNA or targeted siRNA (Dharmacon ON-TARGETplus SMARTpool siRNA) in Opti-MEM reduced-serum medium (Gibco) was transfected at a final concentration of 50 nM with Lipofectamine RNAiMAX (Life Technologies). After 18 hours, the medium was changed to fresh BMDM differentiation medium and incubated for an additional 54 hours before use.

### Cell counting

Macrophages were seeded at 100,000/well in a 12-well plate in 10% HI-FBS DMEM. Macrophages that were treated with ACs were chased for 24 hours with or without various inhibitors. For experiments with M-CSF, macrophages were treated with 50 ng/mL M-CSF (Peprotech) with or without the inhibitors for 24 hours. Cells were detached with 10 mM EDTA in PBS on ice by gentle scraping. Collected cells were counted using a Countess II Automated Cell Counter (Invitrogen).

### Immunoblotting

BMDMs were lysed in 2x Laemmli sample buffer (Bio-Rad) with 50 mM β-mercaptoethanol (Sigma) and lysates were boiled at 95-100°C for 5 minutes. Lysates were loaded onto 4-20% SDS-PAGE gradient gels (Invitrogen) and run at 120V for 90 minutes. Protein was transferred onto a 0.45-micron nitrocellulose membrane at 250 mA for 100 minutes. Membranes were blocked for 1 hour with 5% skim milk in Tris-buffered saline with Tween-20 (TBST). Membranes were then incubated overnight at 4°C with primary antibody diluted in 5% skim milk in TBST. Membranes were washed 3 x 5 minutes with TBST before incubation with HRP-conjugated secondary antibodies at room temperature for 1-2 hours and washed 3 x 5 minutes with TBST. The following antibodies were used: rabbit anti-SIRT1 (CST 9475; 1:1000), rabbit anti-LDHA (ProteinTech 19987-1-AP; 1:1000), rabbit anti-pAMPK (CST 2535; 1:1000), rabbit anti-AMPK (CST 5831; 1:1000), rabbit anti-pCREB (CST 9198; 1:1000), rabbit anti-CREB (CST 4820; 1:1000), rabbit anti-acetyl-cMyc (Sigma ABE26; 1:1000), rabbit anti-cMyc (CST 18583; 1:1000), anti-β-actin HRP-conjugate (CST 5125; 1:10,000), and anti-rabbit IgG HRP-linked (CST 7074; 1:2500). Blots were imaged with film and quantified by ImageJ.

### Immunofluorescence staining and microscopy

BMDMs that were transfected with scrambled or Ldha siRNA were seeded on 8-well chamber slides at 60,000 cells per well before incubation with PKH26-labeled ACs and chased for 3 hours. The cells were fixed for 10 minutes at room temperature in 4% paraformaldehyde. Fixed cells were rinsed with PBS and then permeabilized with 0.25% Triton X-100 for 10 minutes at room temperature, blocked with 2% BSA in PBS for 1 hour at room temperature, and incubated with Myc primary antibody (CST 18583; 1:100) in blocking solution at 4°C overnight. The cells were then rinsed with PBS and incubated with chicken anti-rabbit AF647 secondary antibody (Invitrogen; A-21443) for 1 hour at room temperature in the dark. Cells were rinsed again with PBS and then stained with Hoechst-33342 (CST 4082; 1:1000) for 15 minutes at room temperature before mounting and imaging. For paraffin-embedded tissues, sections were deparaffinized with xylene, rehydrated with 100% ethanol and then 70% ethanol, and rinsed with PBS. Antigen retrieval was performed using 1x Citrate-Based Antigen Unmasking Solution (Vector) and pressure cooking for 10 minutes. TUNEL staining was conducted following antigen retrieval using the *In Situ* Cell Death Detection Kit (Sigma) according to the manufacturer’s instructions. The tissues were blocked with 2% BSA in PBS for 1 hour at room temperature and incubated overnight at 4°C with primary antibody in blocking solution. The following primary antibodies were used: rabbit anti-cMyc (CST 18583; 1:100), rabbit anti-LDHA (ProteinTech 19987-1-AP; 1:100), rabbit anti-Ki67 (ab16667; 1:200), and rat anti-Mac2 (Cedarlane CL8942LE; 1:500). Tissues were rinsed with PBS and incubated for 2 hours at room temperature with fluorescently labeled secondary antibody diluted in blocking solution. The following secondary antibodies were used: goat anti-rabbit AF488 (Invitrogen A11034; 1:200), and goat anti-rat AF647 (Invitrogen A21247; 1:200). All imaging was performed using a Leica epifluorescence microscope (DMI6000B).

### NAD^+^:NADH ratio assay

NAD^+^:NADH ratio was quantified using an NAD/NADH colorimetric assay kit (Abcam). Briefly, WT BMDMs, *Ldha^fl/fl^* BMDMs, or *Ldha^fl/fl^*; *LysMCre*^+/-^ BMDMs were seeded at 2,000,000/well in 6-well plates and incubated with unlabelled ACs, washed with PBS, and chased for 1 hour with or without 10 μM Compound C (Sigma) or 10 mM lactic acid (Sigma). The cells were collected with 400 μL NADH/NAD extraction buffer from the kit, and the assay was carried out according to the manufacturer’s instructions.

### SIRT1 activity assay

SIRT1 activity was measured using a SIRT1 activity assay kit (Abcam) according to the manufacturer’s instructions. Briefly, WT BMDMs were seeded at 4,000,000/dish in 100mm dishes and incubated with unlabelled ACs, washed with PBS, and chased for 1 hour with or without 50 μM FX11 (Sigma), 10 mM lactic acid (Sigma), or 10 μM Compound C (Sigma). Nuclear fractions were isolated and assessed for SIRT1 activity by measuring the fluorescence with an excitation wavelength of 350 nm and emission wavelength of 450 nm.

### Efferocytosis assay

Macrophages were incubated with PKH26-labelled ACs for 45 minutes and then rinsed with PBS to remove unbound ACs and fixed with 4% PFA for 10 minutes at room temperature. After rinsing again with PBS, the macrophages were subjected to brightfield and fluorescence imaging using a Leica DMI6000B inverted epifluorescence microscope to identify macrophages that had taken up a fluorescent AC. AC uptake was quantified as the percent of AC-positive macrophages out of the total number of macrophages per field of view.

### Reverse transcription-quantitative polymerase chain reaction (RT-qPCR)

RNA was isolated from macrophages using the PureLink RNA Mini kit (Life Technologies) according to the manufacturer’s protocol. The RNA quality and concentration was measured using a NanoDrop spectrophotometer (ThermoFisher). cDNA synthesis was generated from RNA using oligo(dT) and Superscript II (Applied Biosystems). RT-qPCR was conducted using the 7500 Real-Time PCR system (Applied Biosystems) and the Power SYBR Green PCR Master Mix (Applied Biosystems). The primers are listed in Supplementary Table 1. Expression of genes of interest were normalized to the expression of the housekeeping gene *Hprt.*

### Statistical analyses

GraphPad Prism software (v.9.4.0) was used for all statistical analyses. Data were tested for normality using the Shapiro-Wilk test. Data that passed the normality test were analyzed using the two-tailed Student’s t-test for comparison of two groups or one- or two-way analysis of variance (ANOVA) with Bonferroni post-hoc analysis for comparison of more than two groups. Data are shown as mean values ± SEM, and differences were considered statistically significant at p < 0.05. For normalization of data, all values were divided by the mean of the control group. The number of mice used for the dexamethasone-thymus assay was determined based on power calculations, with an expected 15-25% coefficient of variation and an 80% chance of detecting a 33% difference in key parameters including in situ efferocytosis and necrotic area.

## Results

### LDHA-dependent lactate production during efferocytosis increases Myc protein and is necessary for EIMP

To test the role of efferocytosis-induced lactate (EIL) in efferocytosis-induced macrophage proliferation (EIMP), we inhibited LDHA with FX11^22^ in efferocytosing murine bone marrow-derived macrophages (BMDMs) and human monocyte-derived macrophages (HMDMs). We first confirmed that efferocytosis increases lactate production and that lactate levels are lowered by treatment with the LDHA inhibitor FX11 (**Extended Data Fig. 1a**). As previously reported^10^, incubating macrophages with ACs to trigger efferocytosis results in an increase in cell number after 24 hours, and we found that this increase was blocked by FX11 and restored by adding back lactic acid to the FX11-inhibited macrophages (**Fig. 1a,b**). In the context of our hypothesis that EIL might stabilize Myc protein to promote EIMP (below), we assessed whether EIL promotes AC-induced Myc protein expression. We employed three methods of blocking LDHA activity: siLdha-treated BMDMs, which led to ~65% silencing of *Ldha* (**Extended Data Fig. 1b**); FX11-treated BMDMs; and BMDMs from *Ldha^fl/fl^*; *LysMCre*^+/-^ (M-LDHA-KO) mice. All three strategies resulted in an attenuation of AC-induced Myc protein expression (**Fig. 1c-e**). FX11 did not affect primary AC engulfment (**Extended Data Fig. 1c**), indicating that the decrease in Myc in LDHA-deficient or LDHA-inhibited macrophages was not merely due to decreased AC uptake. Furthermore, loss of AC-induced Myc protein expression in LDHA KO macrophages could be rescued by exogenous addition of lactic acid (**Fig. 1f**). The link between lactate and EIMP was through Myc, as the decrease in EIMP that occurs with Myc silencing could not be rescued by exogenous lactic acid (**Fig. 1g and Extended Data Fig. 1d**). Moreover, lactic acid treatment in the absence of ACs had no effect on basal Myc protein expression (**Extended Data Fig. 1e**), indicating the importance of first increasing *Myc* mRNA through the efferocytosis-induced p-ERK and AC-nucleotide/Dnase2a/DNA-PK pathways^10^. Together, these data show that EIL is needed to increase Myc protein in efferocytosing macrophages to drive EIMP.

**Fig. 1.**
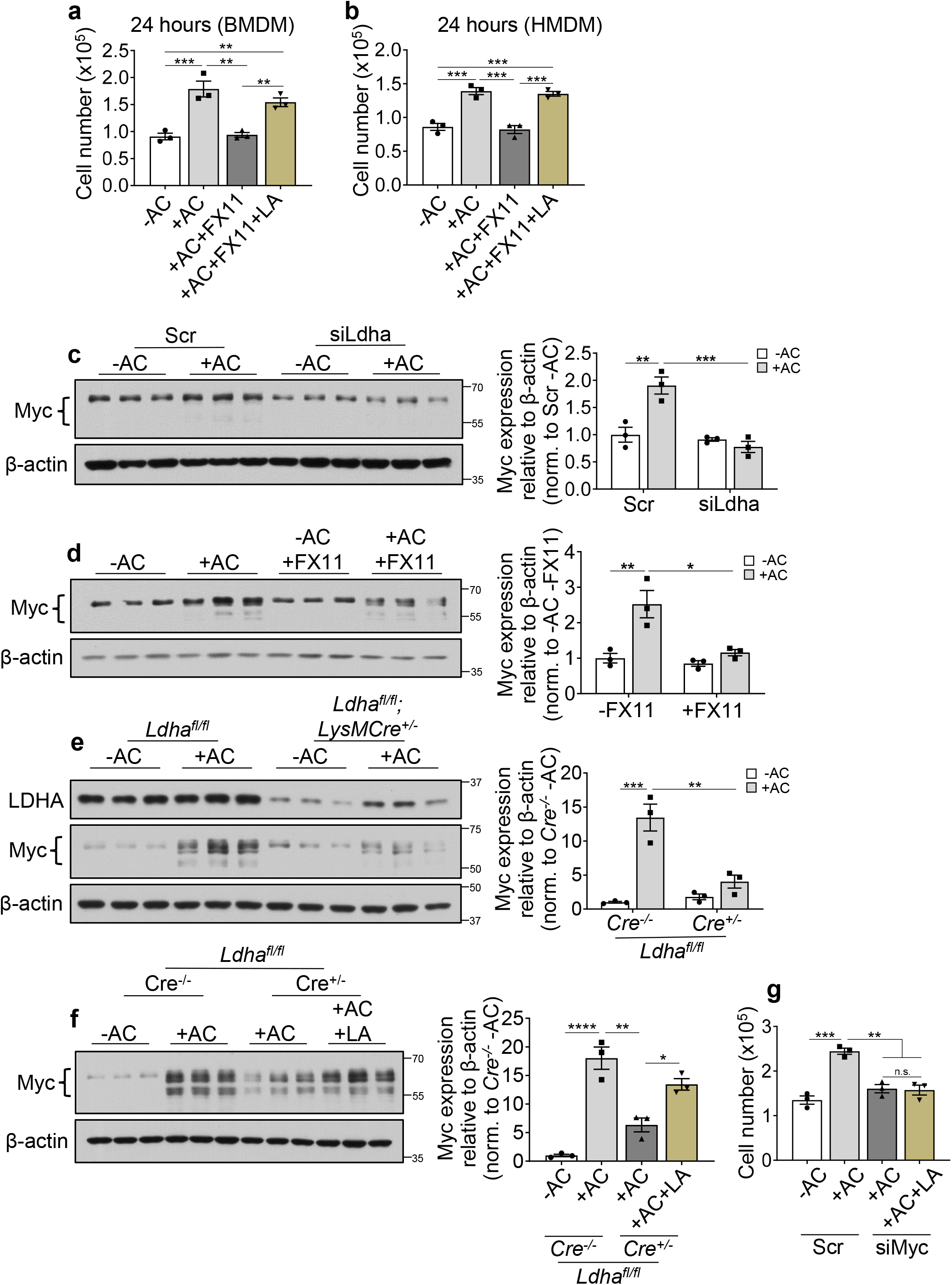
LDHA-dependent lactate production during efferocytosis increases Myc protein and promotes EIMP. **a,b,** BMDMs or HMDMs were incubated with or without ACs for 45 minutes, chased for 24 hours + 50 μM FX11 and + 10 mM lactic acid (LA), and quantified for cell number (n=3). **c,** BMDMs transfected with 50 nM scrambled or Ldha siRNA for 72 hours were incubated with or without ACs for 45 minutes, chased for 3 hours, and immunoblotted for Myc (n=3). **d,** BMDMs were incubated with or without ACs for 45 minutes, chased for 3 hours + 50 μM FX11, and immunoblotted for Myc (n=3). **e,** *Ldha^fl/fl^* or *Ldha^fl/fl^*; *LysMCre*^+/-^ BMDMs were incubated with or without ACs for 45 minutes, chased for 3 hours, and immunoblotted for Myc (n=3). **f,** *Ldha^fl/fl^* or *Ldha^fl/fl^*; *LysMCre*^+/-^ BMDMs were incubated with or without ACs for 45 minutes, chased for 3 hours + 10 mM LA, and immunoblotted for Myc (n=3). **g,** BMDMs transfected with 50 nM scrambled or Myc siRNA were incubated with or without ACs for 45 minutes, chased for 24 hours + 10 mM LA, and quantified for cell number (n=3). Bars represent means + SEM. Statistics were performed by one-way ANOVA in panels a, b, and f-g or two-way ANOVA in panels c-e. *\*P* < 0.05, \*\**P* < 0.01, \*\*\**P* < 0.001, ****P < 0.0001.

### EIL stabilizes Myc protein through Myc deacetylation following AC-nucleotide/DNase2a-mediated induction of *Myc* mRNA

We propose that EIL increases Myc protein in a step that follows and requires *Myc* mRNA induction by the AC-nucleotide/Dnase2a/DNA-PK pathway^10^. If so, then lactic acid should not be able to rescue EIMP if this initial *Myc* mRNA-inducing pathway is blocked. One way to block the AC-nucleotide-DNA-PK pathway is to silence DNase2a, as DNase2a-mediated hydrolysis of AC-DNA to oligonucleotides in phagolysosomes is required to activate DNA-PK and induce *Myc*^10^. Accordingly, we tested the effect of lactic acid on Myc in siDnase2a-treated efferocytosing macrophages. As expected^10^, siDnase2a treatment of efferocytosing macrophages blunted AC-induced Myc protein expression, and, importantly, Myc protein expression was not rescued by lactic acid in the siDnase2a-treated macrophages (**Fig. 2a**). This latter finding is consistent with the idea that lactate stabilizes Myc protein following DNase2a-dependent induction of *Myc* mRNA in efferocytosing macrophages. Further, if lactate promotes EIMP by promoting Myc protein stabilization and not *Myc* transcription, then blocking EIL should not lower AC-induced *Myc* mRNA. Indeed, in both BMDMs and HMDMs, the AC-induced increase in *Myc* mRNA was not blocked by FX11 (**Fig. 2b,c**). In a similar vein, while lactate secreted from efferocytosing macrophages could act on neighboring non-efferocytosing macrophages, these macrophages should not respond with increased Myc protein, as non-efferocytosing macrophages lack the initial AC-induced increase in *Myc* mRNA. Consistent with this prediction, Myc protein expression was increased in AC^+^ but not AC^-^ macrophages, and it was reduced by siLdha only in AC^+^ macrophages (**Fig. 2d**).

**Fig. 2.**
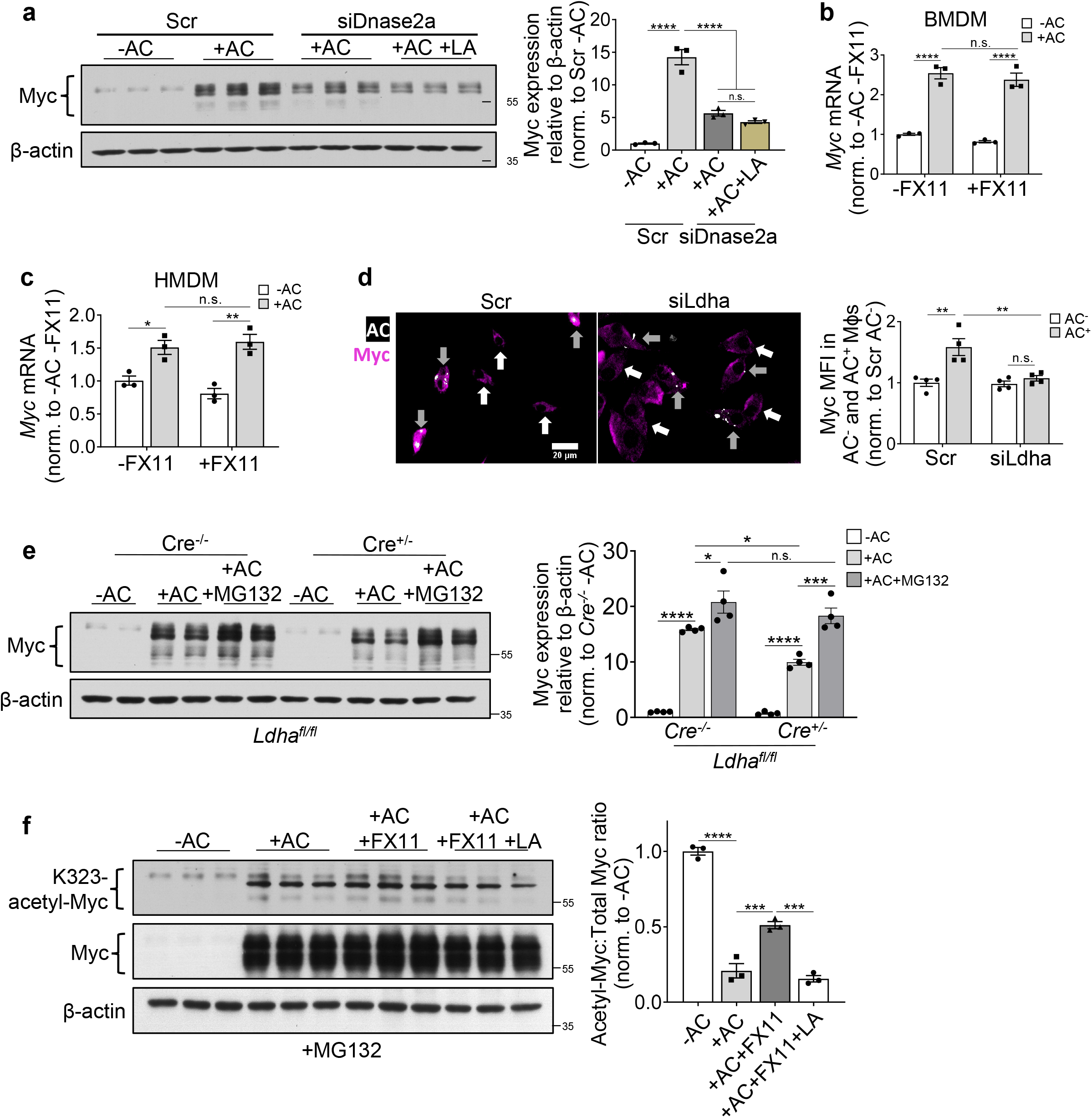
EIL stabilizes AC-induced Myc protein through decreased Myc acetylation, which occurs following AC-nucleotide/DNase2a-mediated *Myc* transcription. **a,** BMDMs transfected with 50 nM scrambled or *Dnase2a* siRNA for 72 hours were incubated with or without ACs for 45 minutes, chased for 3 hours, and immunoblotted for Myc (n=3). **b,c,** BMDMs or HMDMs were incubated with or without ACs for 45 minutes, chased for 3 hours + 50 μM FX11, and assayed for *Myc* mRNA (n=3). **d**, BMDMs were incubated with PKH26-labelled ACs for 45 minutes, chased for 3 hours + 50 μM FX11, fixed with 4% PFA, and immunostained for Myc. Cells were imaged with a 20x objective. Scale bar, 50 μm. **e,** *Ldha^fl/fl^* or *Ldha^fl/fl^*; *LysMCre*^+/-^ BMDMs were incubated with or without ACs for 45 minutes, chased for 3 hours + 10 μM MG132, and immunoblotted for Myc (n=4). **f,** BMDMs were incubated with or without ACs for 45 minutes, chased for 3 hours + 50 μM FX11 and + 10 mM LA in the presence of 10 μM MG-132, and immunoblotted for K^323^-acetyl-Myc and total Myc (n=3). Bars represent means + SEM. Statistics were performed by one-way ANOVA in panels a and f or two-way ANOVA in panels b-e. *\*P* < 0.05, *\*\*P* < 0.01, *\*\*\*P* < 0.001, *\*\*\*\*P* < 0.0001.

Acetylation of Myc promotes its proteasomal degradation^14,15,17^. We therefore first determined if the loss of AC-induced Myc protein expression resulting from LDHA KO could be rescued by the proteasome inhibitor MG-132. In control *Ldha^fl/fl^*) BMDMs, MG-132 treatment boosted AC-induced Myc protein expression, suggesting that there is a basal level of Myc degradation in efferocytosing macrophages (**Fig. 2e, Cre^-/-^ data**). Most importantly, the reduction in AC-induced Myc protein expression in LDHA-KO BMDMs was increased by MG-132 to a level similar to that in control macrophages (**Fig. 2e, Cre^+/-^ data**), These data suggest that EIL maintains the increase in AC-induced Myc protein expression by reducing Myc protein proteasomal degradation. Acetylation at lysine-323 (Ac-K323) of Myc enhances Myc proteasomal degradation^16^. As expected, total Myc protein was very low in baseline macrophages, but acetyl-Myc was present, resulting in a relatively high acetyl-Myc: total Myc ratio (**Fig. 2f, 1^st^ triplicate**). Upon incubation with ACs, both acetyl-Myc and total Myc increased, but the increase in total Myc was much greater, resulting in a marked lowering of the acetyl-Myc: total Myc ratio (**Fig. 2f, 2^nd^ triplicate**). As predicted, FX11 did not lower total Myc owing to the presence of MG-132, but it did raise acetyl-Myc, thereby increasing the ratio (**Fig. 2f, 3^rd^ triplicate**), and this increase was abrogated by adding back lactic acid (**Fig. 2f, 4^th^ triplicate**). These data, when combined with those above, support the idea that after *Myc* transcription is induced by the AC-cargo pathway^10^, EIL deacetylates and stabilizes Myc protein to enable macrophage proliferation.

### EIL-induced SIRT1 activation is necessary for increased Myc protein and EIMP

Sirtuin-1 (SIRT1) is an NAD^+^-dependent protein deacetylase that has been shown previously in other cell types to deacetylate and stabilize Myc protein^14–16,18^, and SIRT1 has also been implicated in efferocytosis and inflammation resolution^12,23^. Moreover, lactic acid generated by exercise has been reported to activate SIRT1 in the brain^24^. We therefore considered the hypothesis that EIL uses SIRT1 to stabilize Myc protein and enable EIMP. In support of this idea, we found that efferocytosis activates SIRT1, which was attenuated by FX11 treatment and restored with exogenous lactic acid (**Fig. 3a**). Efferocytosis also increased SIRT1 protein expression, and partial silencing of SIRT1 lowered AC-induced Myc protein expression (**Fig. 3b**). SIRT1-silencing caused a modest decrease in primary efferocytosis (**Extended Data Fig. 1f)**, which was likely too small to explain the large decrease in AC-induced Myc protein expression. Nonetheless, we tested the effect of the SIRT1 inhibitor EX527^25^ added after AC uptake and found that this treatment also lowered AC-induced Myc protein expression (**Fig. 3c**). Moreover, exogenous lactic acid could not rescue the reduction in AC-induced Myc protein expression following EX527 treatment (**Extended Data Fig. 1g**), which is consistent with SIRT1 acting downstream of lactate. In addition, EX527 did not affect AC-induced *Myc* mRNA expression, indicating that SIRT1 regulates Myc protein expression post-transcriptionally (**Fig. 3d)**. Most importantly, SIRT1 inhibition blocked EIMP, as measured by total cell number (**Fig. 3e**). To evaluate the specificity of the lactate-SIRT1 pathway for macrophage proliferation triggered by efferocytosis, we tested whether FX11 and EX527 affect CSF1-induced macrophage proliferation, a common form of proliferation observed in inflammation that is distinct from EIMP^10,26^. We found that neither inhibitor blocked the increase in cell number in this setting (**Fig. 3f**). Finally, to directly link SIRT1 to stabilization of Myc against proteasomal degradation (above), we showed that the reduction in AC-induced Myc protein expression by EX527 was abrogated by MG-132 (**Fig. 3g**). Furthermore, EX527 increased the acetyl-Myc: total Myc protein ratio in efferocytosing macrophages (**Fig. 3h**). These combined data support the idea that EIL-induced SIRT1 activation deacetylates and stabilizes Myc protein, thereby enabling EIMP.

**Fig. 3.**
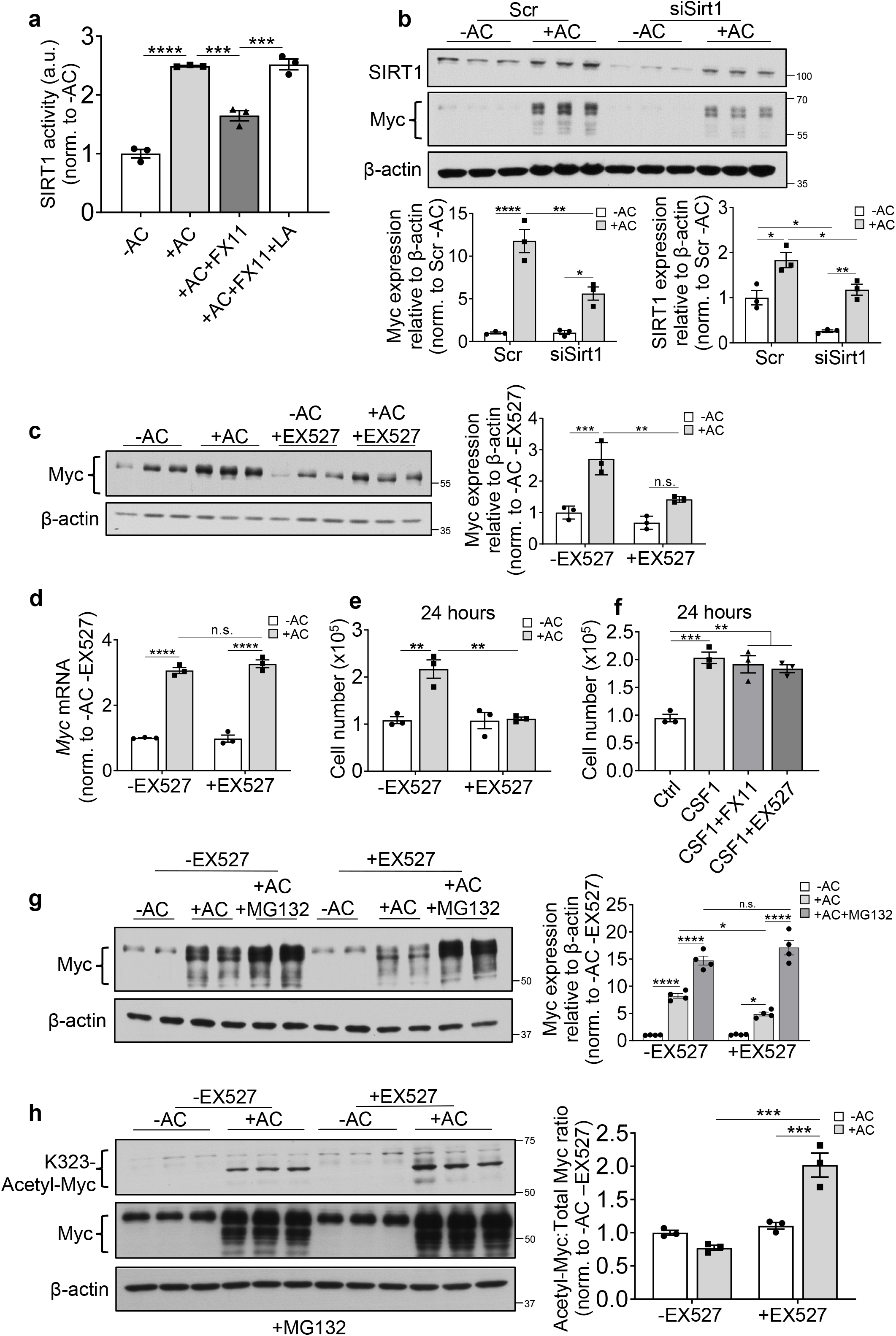
EIL activates SIRT1, which stabilizes Myc by deacetylation and promotes EIMP. **a,** BMDMs were incubated with or without ACs for 45 minutes, chased for 1 hour + 50 μM FX11 with or without 10 mM LA, and assayed for SIRT1 activity (n=3). **b,** BMDMs transfected with 50 nM scrambled or *Sirtl* siRNA for 72 hours were incubated with or without ACs for 45 minutes, chased for 3 hours, and immunoblotted for SIRT1 and Myc (n=3). **c,** BMDMs were incubated with or without ACs for 45 minutes, chased for 3 hours + 10 μM EX527, and immunoblotted for Myc (n=3). **d,** BMDMs were incubated with or without ACs for 45 minutes, chased for 3 hours + 10 μM EX527, and assayed for *Myc* mRNA (n=3). **e,** BMDMs were incubated with or without ACs for 45 minutes, chased for 24 hours + 10 μM EX527, and quantified for cell number (n=3). **f,** BMDMs were incubated with or without ACs for 45 minutes, treated for 24 hours with 50 ng/mL CSF-1 + 50 μM FX11 or 10 μM EX527, and quantified for cell number (n=3). **g,** BMDMs were incubated with or without ACs for 45 minutes, chased for 3 hours + 10 μM EX527 and + 10 μM MG132, and immunoblotted for Myc (n=4). **h)** BMDMs were incubated with or without ACs for 45 minutes, chased for 3 hours + 10 μM EX527 in the presence of 10 μM MG-132, and immunoblotted for K^323^-acetyl-Myc and total Myc (n=3). Bars represent means + SEM. Statistics were performed by one-way ANOVA in panels a and f or two-way ANOVA in panels b-e and g-h. \**P* < 0.05, \*\**P* < 0.01, \*\*\**P* < 0.001, \*\*\*\**P* < 0.0001, n.s. = no significance.

### EIL activates AMPK, thereby increasing NAD^+^:NADH ratio, SIRT1 activity, Myc protein expression, and EIMP

Several previous studies on lactate and SIRT1, although not related to Myc or proliferation, suggested to us a hypothesis as to how lactate might be linked to the SIRT1-Myc-EIMP pathway. SIRT1 is an NAD^+^-dependent protein deacetylase^27^. In non-immune tissues like skeletal muscle, AMPK has been shown to raise NAD^+^ levels through the nicotinamide phosphoribosyltransferase (NAMPT) salvage pathway, which then activates SIRT1^27,28^. Other studies have reported that lactic acid can activate AMPK, e.g., in skeletal muscle *in vivo* and BMDMs *in vitro*^7,29^, and that efferocytosis can activate AMPK^30^. Given these previous observations, we hypothesized that EIL might activate AMPK, which would then increase the NAD^+^:NADH ratio and activate the SIRT1-mediated Myc-EIMP pathway. Consistent with this hypothesis, we found that both lactic acid treatment and incubation of macrophages with ACs raised the level of pAMPK, which indicates AMPK activation, and the lactic acid-induced increase in pAMPK in efferocytosing macrophages was prevented by LDHA silencing (**Fig. 4a,b**). Moreover, inhibition of AMPK with the inhibitor Compound C reduced AC-induced SIRT1 activity (**Fig. 4c)**. Compound C also reduced AC-induced increases in both Myc protein (**Fig. 4d**) and EIMP, as measured by cell count (**Fig. 4e**). These effects were not caused by a decrease in AC uptake, as AMPK inhibition by Compound C did not affect primary efferocytosis (**Extended Data Fig. 1h**). Furthermore, AMPK did not blunt CSF1-dependent macrophage proliferation, indicating specificity for EIMP (**Fig. 4f**).

**Fig. 4.**
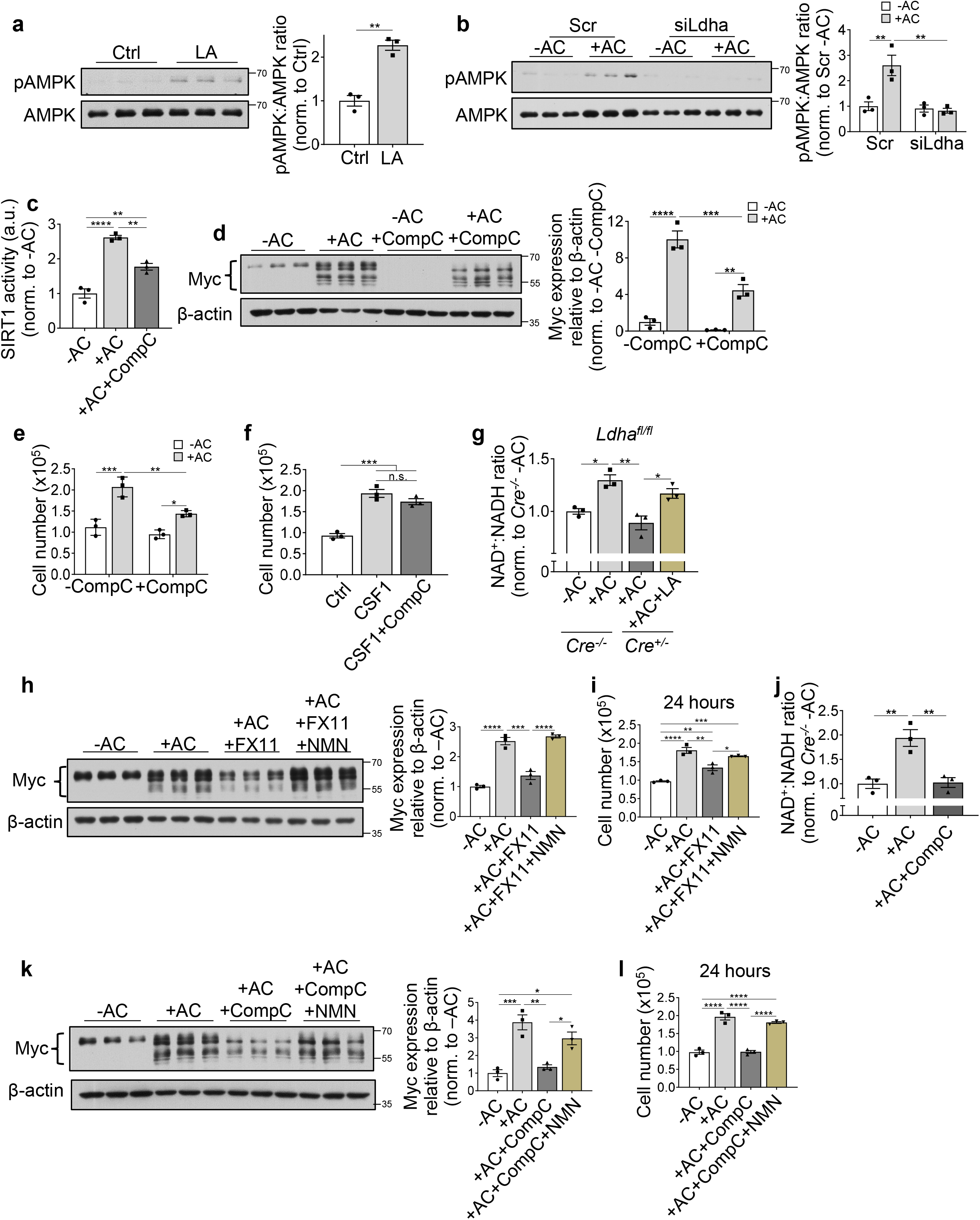
EIL activates AMPK, thereby increasing the NAD^+^:NADH ratio, SIRT1 activity, Myc protein, and EIMP. **a,** BMDMs were incubated with or without ACs for 45 minutes, treated + 10 mM LA for 1 hour, and immunoblotted for pAMPK and total AMPK (n=3). **b,** BMDMs transfected with 50 nM scrambled or Ldha siRNA for 72 hours were incubated with or without ACs for 45 minutes, chased for 1 hour, and immunoblotted for pAMPK and total AMPK (n=3). **c,** BMDMs were incubated with or without ACs for 45 minutes, chased for 1 hour + 10 μM CompC, and assayed for SIRT1 activity (n=3). **d,** BMDMs were incubated with or without ACs for 45 minutes, chased for 3 hours + 10 μM Compound C (CompC), and immunoblotted for Myc (n=3). **e,** BMDMs were incubated with or without ACs for 45 minutes, chased for 24 hours + 10 μM CompC, and quantified for cell number (n=3). **f,** WT BMDMs were treated for 24 hours with 50 ng/mL CSF-1 + 10 μM CompC and then quantified for cell number (n=3). **g,** *Ldha^fl/fl^* or *Ldha^fl/fl^*; *LysMCre*^+/-^ BMDMs were incubated with or without ACs for 45 minutes, chased for 1 hour + 10 mM LA, and assayed for NAD^+^:NADH ratio (n=3). **h,** BMDMs were incubated with or without ACs for 45 minutes, chased for 3 hours + 50 μM FX11 and + 500 μM nicotinamide mononucleotide (NMN), and immunoblotted for Myc (n=3); or **i,** chased for 24 hours + 50 μM FX11 and + 500 μM NMN and quantified for cell number (n=3). **j,** BMDMs were incubated with or without ACs for 45 minutes, chased for 1 hour + 10 μM CompC, and assayed for NAD^+^:NADH ratio (n=3). **k,** BMDMs were incubated with or without ACs for 45 minutes, chased for 3 hours + 10 μM CompC and + 500 μM nicotinamide mononucleotide (NMN), and immunoblotted for Myc (n=3). **l,** BMDMs were incubated with or without ACs for 45 minutes, chased for 24 hours + 10 μM CompC + 500 μM NMN, and quantified for cell number (n=3). Bars represent means + SEM. Statistics were performed by student’s t-test in panel a, one-way ANOVA in panels c, and f-l, or two-way ANOVA in panels d and e. \**P* < 0.05, \*\**P* < 0.01, \*\*\**P* < 0.001, \*\*\*\**P* < 0.0001, n.s. = no significance.

As mentioned, EIL might be linked to SIRT1 activation through an increase in the NAD^+^:NADH ratio^27^. Consistent with this idea, the addition of ACs to macrophages increased the NAD^+^:NADH ratio, and this increase was prevented by LDHA-KO and was rescued by exogenous lactic acid treatment (**Fig. 4g**). To assess causation, FX11-treated macrophages were supplemented with or without the NAD^+^ precursor, nicotinamide mononucleotide (NMN), to rescue NAD^+^ levels and then assayed for Myc protein expression and proliferation. We found that the decreases in both Myc protein expression and cell number in FX11-treated AC-exposed macrophages were rescued by NMN (**Fig. 4h,i**). In addition, Compound C blunted the AC-induced increase in the NAD^+^:NADH ratio (**Fig. 4j**), and NMN supplementation of Compound C-treated cells, which restored the NAD+:NADH ratio (**Extended Data Fig. 1i**), rescued the decreases in both AC-induced Myc protein expression and cell number (**Fig. 4k,l**). In the absence of efferocytosis, NMN did not affect Myc protein expression (**Extended Data Fig. 1j**). Furthermore, NMN in these experiments was acting through SIRT1, as the loss of AC-induced Myc protein in SIRT1-silenced BMDMs could not be rescued by NMN (**Extended Data Fig. 1k**). Thus, EIL activates an AMPK-NAD^+^-SIRT1 pathway to increase Myc protein expression and promote EIMP.

### EIL activates AMPK by GPR132-PKA signaling to increase Myc protein and EIMP

To determine how EIL activates AMPK, we considered the canonical mechanism of AMPK activation, namely, as a response to ATP depletion^31^. However, ATP levels do not change or slightly increase rather than decrease during and after AC uptake^8,12^. We therefore became interested in a cell-surface G-protein coupled lactate receptor called GPR132, which was shown previously to be involved in lactate-mediated activation of a PKA-AMPK pathway in inflammatory macrophages^7^. If this pathway were relevant in our setting, EIL-induced Myc protein would require the release of lactate to the extracellular space to activate the cell-surface receptor. Indeed, partial silencing the lactate transporter, SLC16a1, which prevents the export of lactate from efferocytosing macrophages^8,9^, partially lowered Myc protein expression (**Fig. 5a and Extended Data Fig. 1l**). We next explored the effect of the GPR132 inhibitor telmisartan^32^ and the GPR132 activator ONC212^33^ on EIL-induced Myc protein expression. Telmisartan treatment resulted in a reduction of AC-induced Myc protein expression (**Fig. 5b**), and, conversely, the loss of AC-induced Myc protein expression in FX11-treated macrophages was rescued by ONC212 (**Fig. 5c**). As a complement to the telmisartan data, we showed that siRNA-mediated silencing of GPR132 lowered both AC-induced Myc protein expression and EIMP, as measured by cell number (**Fig. 5d,e and Extended Data Fig. 1m**).

**Fig. 5.**
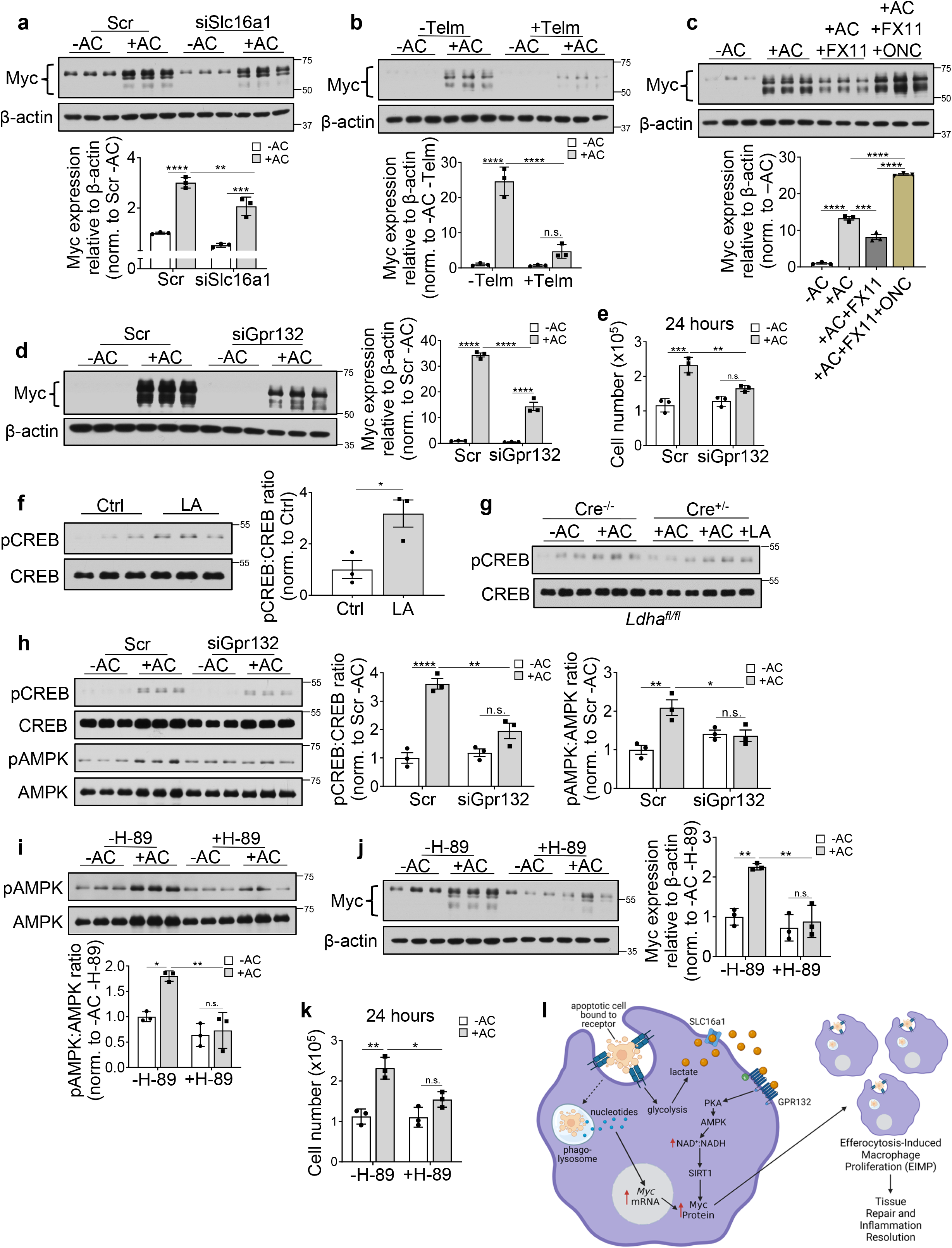
EIL activates AMPK by GPR132-PKA signaling, which increases Myc protein and EIMP. **a,** BMDMs transfected with 50 nM scrambled or *Slc16a1* siRNA for 72 hours were incubated with or without ACs for 45 minutes, chased for 3 hours, and immunoblotted for Myc (n=3). **b,** BMDMs were incubated with or without ACs for 45 minutes, chased for 3 hours + 10 μM telmisartan (Telm), and immunoblotted for Myc (n=3). **c,** BMDMs were incubated with or without ACs for 45 minutes, chased for 3 hours + 50 μM FX11 and + 10 μM ONC212 (ONC), and immunoblotted for Myc (n=3). **d-e,** BMDMs transfected with 50 nM scrambled or Gpr132 siRNA for 72 hours were incubated with or without ACs for 45 minutes, chased for 3 hours, and immunoblotted for Myc or quantified for cell number (n=3). **f,** BMDMs were treated + 10 mM LA for 1 hour and then immunoblotted for pCREB and total CREB (n=3). **g,** *Ldha^fl/fl^* or *Ldha^fl/fl^*; *LysMCre*^+/-^ BMDMs were incubated with or without ACs for 45 minutes, chased for 1 hour + 10 mM LA, and immunoblotted for pCREB and total CREB (n=3). **h,** BMDMs transfected with scrambled or *Gpr132* siRNA were incubated with or without ACs for 45 minutes, chased for 1 hour, and immunoblotted for pCREB, total CREB, pAMPK, and total AMPK (n=3). **i,** BMDMs were incubated with or without ACs for 45 minutes, chased for 1 hour + 10 μM H-89 and immunoblotted for pAMPK and total AMPK (n=3); or **j,** chased for 3 hours + 10 μM H-89 and immunoblotted for Myc (n=3); or **k,** chased for 24 hours + 10 μM H-89 and quantified for cell number (n=3). Bars represent means + SEM. Statistics were performed by student’s t-test in panel f, one-way ANOVA in panels c and g, or two-way ANOVA in panels a-b, d-e, and h-k. \**P* < 0.05, \*\**P* < 0.01, \*\*\**P* < 0.001, \*\*\*\**P* < 0.0001, n.s. = no significance. **l,** Graphical depiction of how lactate produced by efferocytosis-induced macrophage glycolysis (EIMG)^9^ integrates with the previously elucidated AC-derived nucleotide pathway^10^ to enhance Myc protein levels and EIMP.

To probe the role of the GPR132 signaling molecule PKA, we first incubated macrophages with exogenous lactic acid and showed that this treatment increased phosphorylation of CREB (**Fig. 5f**), which is a target of PKA and a marker for PKA activity. Further, incubation of macrophages with ACs increased pCREB, and this increase was reduced by LDHA KO and restored by exogenous lactic acid (**Fig. 5g**). In addition, silencing GPR132 in AC-exposed macrophages lowered pCREB and p-AMPK (**Fig. 5h**). Finally, treatment of AC-exposed macrophages with the PKA inhibitor H-89 reduced pAMPK activation, Myc protein expression, and EIMP, as measured by cell count (**Fig. 5i-k**).

In conclusion, while our previous study elucidated an AC-cargo pathway in efferocytosing macrophages that increases *Myc* mRNA^10^, our new data suggest that EIMP requires a second, post-transcriptional process, namely, stabilization of Myc protein. This process is triggered by extracellular lactate resulting from efferocytosis-induced macrophage glycolysis and lactate export by SLC16a1^8,9^. Extracellular lactate activates a GPR132-PKA-AMPK pathway that increases the NAD^+^:NADH ratio, leading to SIRT1-mediated Myc protein deacetylation and stabilization. The increase in Myc protein from the combined actions of the AC-cargo pathway and EIL enable EIMP (**Fig. 5l**).

### *In vivo* evidence that EIL promotes EIMP and tissue resolution

To test the role of EIL in EIMP and efferocytosis *in vivo,* we utilized a model of acute apoptosis in which efferocytosis is required to promote tissue resolution. In this model, dexamethasone treatment of mice induces thymocyte apoptosis, followed by clearance of the apoptotic thymocytes by recruited thymic macrophages and then tissue resolution^10,11,34, 35 36^. We conducted this experiment in irradiated mice transplanted with either control *Ldha^fl/fl^* bone marrow cells or *Ldha^fl/fl^* bone marrow cells transfected ex vivo with cell-permeable TAT-Cre before infusion to knockdown *Ldha* (LDHA-KD). In the LDHA-KD cohort, LDHA protein was decreased by ~50% in both bone marrow cells (**Fig. 6a**) and thymic macrophages, but not in non-macrophage Mac2^-^ thymic cells (**Fig. 6b and Extended Data Fig. 2a**). Moreover, the concentrations of immune cells in the circulation were similar in the LDHA KD and control group mice, suggesting that hematopoietic LDHA-KD did not affect the systemic immune system (**Extended Data Fig. 2b-g**). Next, we assayed Myc protein expression and EIMP in thymic macrophages by immunostaining for Myc protein and Ki67, a marker or proliferation, respectively. We looked at both macrophages carrying out efferocytosis, as marked by TUNEL^+^ staining in the macrophage cytoplasm (AC^+^ macrophages), as well as macrophages without cytoplasmic TUNEL staining (AC^-^ macrophages). In control mice, there was a marked increase in both Myc and Ki67 in AC^+^ thymic macrophages, but not AC^-^ macrophages, as previously described^10^ (**Fig. 6c,d, Ctrl**). However, both Myc protein expression and Ki67-positivity were substantially less in AC^+^ macrophages in the thymi of LDHA-KD mice (**Fig. 6c,d. LDHA-KD**). Continual efferocytosis is a critical process in the dexamethasone model due to the large number of apoptotic thymocytes^10,36,37^. In this context, we predicted that LDHA-KD, by preventing the pool of efferocytosis-capable macrophages from expanding, would show a defect in AC clearance. In line with this prediction, thymic macrophage efferocytosis, quantified as the ratio of macrophage-associated-to-free TUNEL^+^ dead cells^10,11,34,35^, was impaired in the LDHA-KD cohort (**Fig. 6e**). Further, consistent with impaired dead cell clearance, the thymi of LDHA-KD mice had more TUNEL^+^ cells than control mouse thymi (**Fig. 6f**). Finally, a key functional endpoint of efferocytosis is prevention of tissue necrosis, and we found that thymic necrosis, quantified as the percent area of hypocellular regions with fragmented nuclei, was increased in the LDHA-KD cohort (**Fig. 6g**). These combined data provide evidence that LDHA-dependent EIL is necessary for Myc expression and proliferation in efferocytosing macrophages *in vivo* and, most importantly, for the proper clearance of dead cells and tissue resolution.

**Fig. 6.**
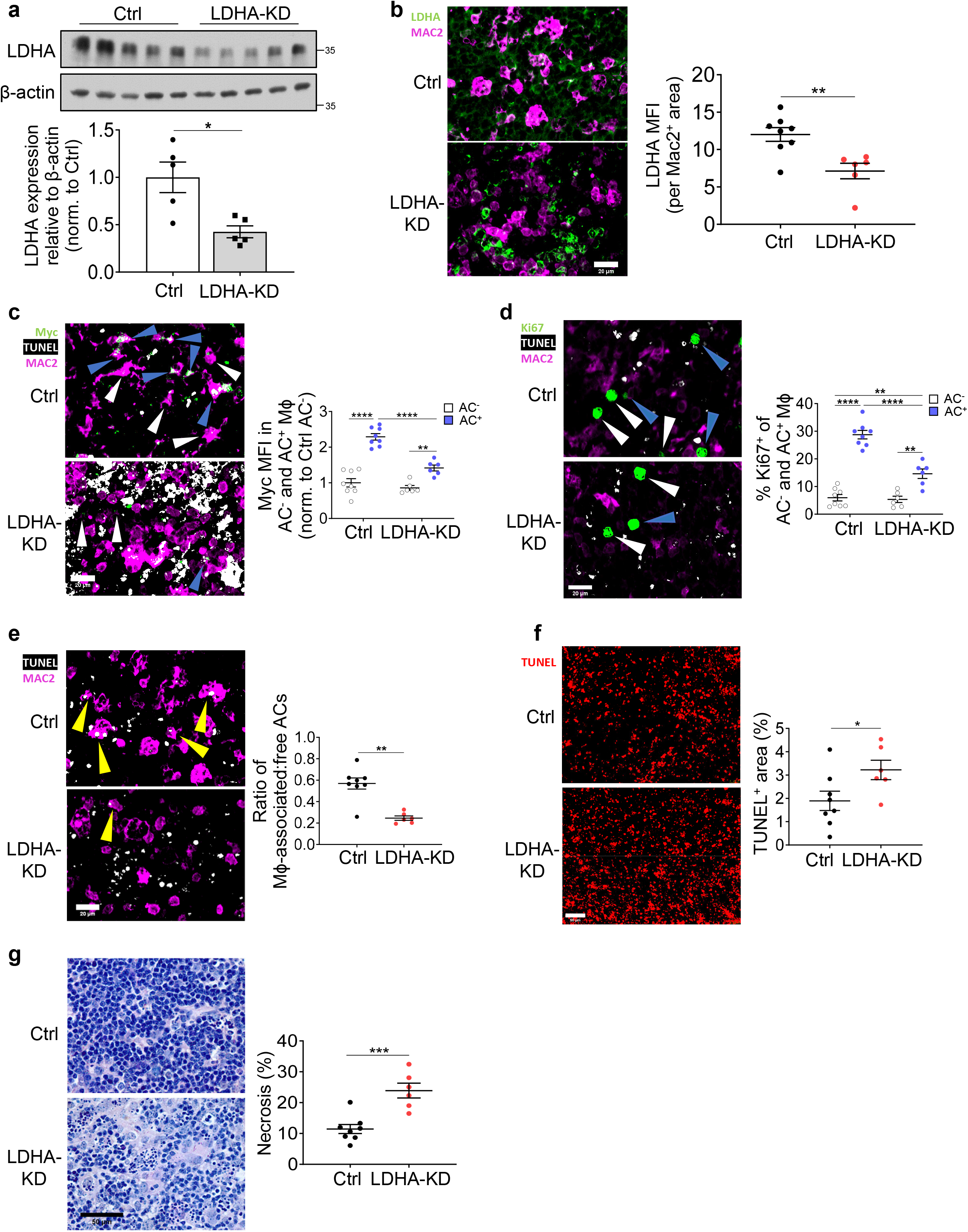
*In vivo* evidence that EIL promotes EIMP and tissue resolution. Bone marrow was isolated from *Ldha^fl/fl^* mice, treated with 5 μM TAT-Cre to knockdown LDHA (LDHA KD) or with vehicle (Ctrl), and transplanted into irradiated male C57BL/6J mice (n=6-8 mice/group). After 4 weeks, the mice were subjected to a dexamethasone-thymus assay. **a,** Bone marrow cells isolated from mice were immunoblotted for LDHA (n=5). **b,** Thymus sections were stained for LDHA (green) and Mac2 (magenta) and quantified for LDHA MFI in Mac2^+^ and Mac2^-^ areas (N-6-8). Scale bar, 20 μm. **c,** Thymus sections stained with TUNEL (white), anti-Myc (green), and anti-Mac2 (magenta) were quantified for Myc MFI in Mac2^+^ TUNEL^-^ (AC^-^) and Mac2^+^ TUNEL^+^ (AC^+^) cells (n=6-8). Scale bar, 20 μm. **d,** Thymus sections stained with TUNEL (white), anti-Ki67 (green) and anti-Mac2 (magenta) were quantified for the % Ki67^+^ Mac2^+^ cells that were TUNEL^-^ (AC^-^) or TUNEL^+^ (AC^+^) (n=6-8). Scale bar, 20 μm. **e,** Thymus sections were stained with TUNEL (white) and anti-Mac2 (magenta), and, as a measure of efferocytosis, the ratio of Mac2^+^ macrophage-associated:free TUNEL^+^ ACs was quantified (n=6-8). Scale bar, 20 μm. **f)** Thymus sections stained with TUNEL (red) were quantified for % TUNEL^+^ area of total area (n=6-8). Scale bar, 50 μm. **g)** H&E-stained thymus sections were quantified for % necrotic area (n=6-8). Scale bar, 50 μm. Dot blots show means + SEM. Statistics were performed by student’s t-test in panels a-b and e-g and by two-way ANOVA in panels c-d. \**P* < 0.05, \*\**P* < 0.01, \*\*\**P* < 0.001, \*\*\*\**P* < 0.0001. n.s., non-significant (*P* > 0.05).

## Discussion

These findings herein illustrate how efferocytosing macrophages can integrate AC-cargo metabolism with macrophage glucose-lactate metabolism to carry out a key process in tissue resolution. In particular, the proliferation of efferocytosing macrophages (EIMP), which expands the pool of pro-resolving macrophages to mediate resolution, requires multiple inputs, consisting not only of MerTK-ERK1/2 and AC nucleotide-DNA-PK-mTORC2-Akt signaling to increase *Myc* mRNA, as described previously^10^, but also efferocytosis-induced lactate (EIL) to subsequently stabilize Myc protein.

Although the pathway described here is mediated by extracellular lactate, which could in theory act on all neighboring macrophages, only AC^+^ macrophages were found to have an LDHA-dependent increase in Myc expression. This finding is consistent with the idea that the EIL-Myc protein stabilization pathway is a process that follows increased *Myc* mRNA, which occurs only in efferocytosing macrophages. However, there may be other paracrine pro-resolving effects of lactate that can affect both AC^+^ and AC^-^ macrophages, including other effects mediated by the GPR132-PKA-AMPK signaling pathway. For example, lactate can drive phenotypic switching of macrophages to a pro-resolving phenotype in certain contexts, including immunosuppression by cancer cells and the resolution of LPS-stimulated macrophage inflammation^6,7,38^, and GPR132 and AMPK signaling have been directly implicated in proresolution phenotypic switching in macrophages^7,38,39^. A previous report showed that conditioned media from efferocytosing macrophages increased *Tgfb* and *Il10* mRNA in non-efferocytosing macrophages and that this did not occur using conditioned media from efferocytosing macrophages whose lactate transporters had been silenced^8^. However, a direct, mechanistic link between EIL and pro-resolving mediators remains to be fully explored. Paracrine effects are important, as only a subset of macrophages carry out efferocytosis^8–10,34^, and thus EIL may be a mechanism by which the resolution-efferocytosis cycle may act more broadly.

Enhanced lactate secretion by efferocytosing macrophages could also have a beneficial role in tissue repair by signaling to other cell types^40,41^. For example, lactate has been shown to promote a synthetic phenotype in vascular smooth muscle cells and enhance their proliferation^20^. In atherosclerosis regression, a setting in which efferocytosis is reawakened after being relatively dormant in progressing lesions^11,42^, a major hallmark is thickening of the fibrous cap mediated by synthetic vascular smooth muscle cell-derived cells, which contributes to plaque stabilization^43^. Accordingly, paracrine effects of EIL could have a role in enhancing this process. Lactate has also been shown to suppress the activation of effector CD4^+^ and CD8^+^ T-cells^44^, whereas lactate promotes the proliferation and activity of T-regulatory (Treg) cells^21^. Treg cells play an important role in polarizing macrophages to a pro-resolving, pro-efferocytic phenotype, including in atherosclerosis regression, and enhance the secretion of factors such as IL-10 by macrophages to dampen inflammation and assist in tissue repair^42,45^. Thus, paracrine effects of EIL may play a role in promoting the efferocytosis-resolution cycle in settings in which Treg cells play important roles, such as atherosclerosis regression.

Both inflammatory macrophages and efferocytic macrophages can undergo proliferation and increase glycolysis to enhance lactate secretion^8–10,26^. However, whereas inflammatory macrophages do not undergo Myc-induced EIMP^10^, they do undergo CSF-1-mediated proliferation^26^. Consistent with these findings, lactate regulates EIMP through stabilizing Myc protein, and lactate is not required for CSF-1-mediated proliferation. Furthermore, GPR132 is known to be downregulated in LPS-induced inflammatory macrophages^7^, which suggests that the downstream effects of lactate are highly context-specific and dictated by the expression of upstream mediators. GPR132 downregulation may be a contributing factor as to why inflammatory macrophages are unable to undergo EIMP despite being able to perform efferocytosis^10^. Interestingly, the addition of exogenous lactate *in vitro* and lactate infusion *in vivo* were shown to increase *Gpr132* mRNA and GPR132 protein^7^, which may form a positive feedback loop and contribute to the beneficial, pro-resolving effects of lactate infusion observed in humans. For example, ischemic postconditioning through the infusion of lactated Ringer’s solution (sodium lactate solution) resulted in an attenuation of ischemia-reperfusion injury in myocardial infarction patients^46^. Exercise also transiently increases circulating lactic acid, which could be a contributing factor in the ability of exercise to protect against chronic inflammatory diseases^47^. From a therapeutic viewpoint, the findings here also raise the possibility that direct activation of GPR132 through pharmacologic agonists such as ONC212 could reawaken the dormant efferocytosis-resolution cycle that drives many chronic inflammatory diseases^2,33^.

## Supporting information

EIL Supplement

## Acknowledgements

This work was supported by an American Heart Association Postdoctoral Fellowship (900337; to M.S.); the Niels Stensen Fellowship (to M.S.); and NIH/NHLBI grants R35-HL145228 and P01-HL087123 (to I.T.). We thank Dr. Lev Becker (University of Chicago) for providing *Ldha^fl/fl^* and *Ldha^fl/fl^*; *LysMCre*^+/-^ mouse femurs for BMDM differentiation to use in our *in vitro* studies. We thank Dr. Xiaobo Wang (Columbia University) for assisting with i.v. injections for bone marrow transplantation. We acknowledge Dr. Caisheng Lu of the Columbia Center for Translational Immunology Core Facility for assisting in the immunofluorescent imaging experiments which were conducted in the Columbia Center for Translational Immunology Core Facility, funded by NIH grants P30CA013696, S10RR027050, and S10OD020056.

## Author contributions

D.N. and I.T. conceived the project. D.N., M.S., and I.T. provided intellectual input to the development of the project. D.N. performed the *in vitro* experiments, and D.N. and M.S. performed the *in vivo* dexamethasone-thymus experiment.

