## Supplementary material for "Efferocytosis-Induced Lactate Enables the Proliferation of Pro-Resolving Macrophages to Mediate Tissue Repair": EIL Supplement

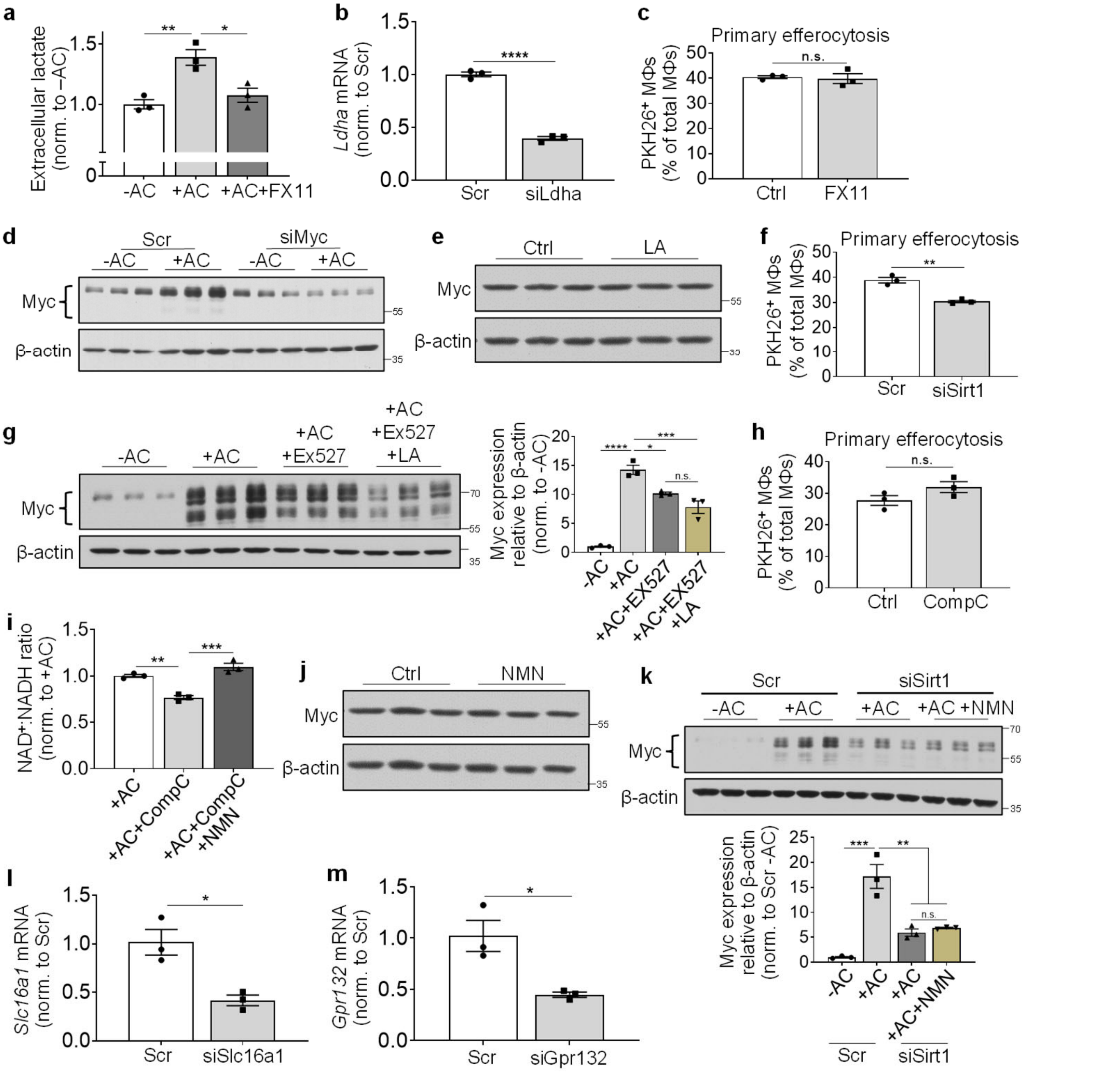


**Extended Data Fig. 1 | Related to Fig. 1-5: Controls for BMDM experiments.**

**a,** BMDMs were incubated with or without ACs for 45 minutes before washing and chasing for 1 hour in low-serum DMEM $\pm$ 50 μM FX11. The media were assayed for lactate (n=3). **b)** BMDMs were transfected with scrambled (Scr) or Ldha siRNA for 72 hours and then assayed for *Ldha* mRNA by RT-qPCR (n=3). **c)** BMDMs pre-treated with 50 μM FX11 for 1 hour were incubated for 45 minutes with PKH26-labelled ACs and quantified for % PKH26$^+$ macrophages (n=3). **d)** BMDMs transfected with 50 nM scrambled or Myc siRNA for 72 hours were incubated with or without ACs for 45 minutes, chased for 3 hours, and immunoblotted for Myc (n=3). **e,** BMDMs were treated $\pm$ 10 mM LA for 3 hours and immunoblotted for Myc (n=3). **f,** BMDMs were transfected with 50 nM scrambled or Sirt1 siRNA for 72 hours, incubated with PKH26-labelled ACs for 45 minutes, and quantified for the percent PKH26$^+$ macrophages (n=3). **g)** BMDMs were chased for 3 hours $\pm$ 10 μM EX527 with or without 10 mM LA and immunoblotted for Myc (n=3). **h)** BMDMs pre-treated for 1 hour with 10 μM CompC were incubated with PKH26-labelled ACs and quantified for the percent PKH26$^+$ (n=3). **i)** BMDMs were chased for 1 hour + 10 μM CompC with or without 500 μM NMN and then assayed for NAD$^+$:NADH ratio (n=3). **j)** BMDMs were treated with 500 μM NMN for 3 hours and then immunoblotted for Myc (n=3). **k)** BMDMs transfected with scrambled or Sirt1 siRNA were chased for 3 hours $\pm$ 500 μM NMN and then immunoblotted for Myc (n=3). **l)** BMDMs were transfected with 50 nM scrambled or Slc16a1 siRNA for 72 hours and then assayed for *Slc16a1* mRNA by RT-qPCR (n=3). **m)** BMDMs were transfected with 50 nM scrambled or Gpr132 siRNA for 72 hours and then assayed for *Gpr132* mRNA by RT-qPCR (n=3). Bars represent means $\pm$ SEM. Statistics were performed by student's t-test in panels b-c, f, h, and l-m or one-way ANOVA in panels a, g, i, and k. *$P$ < 0.05, **$P$ < 0.01, ***$P$ < 0.001, ****$P$ < 0.0001. n.s., non-significant ($P$ > 0.05).

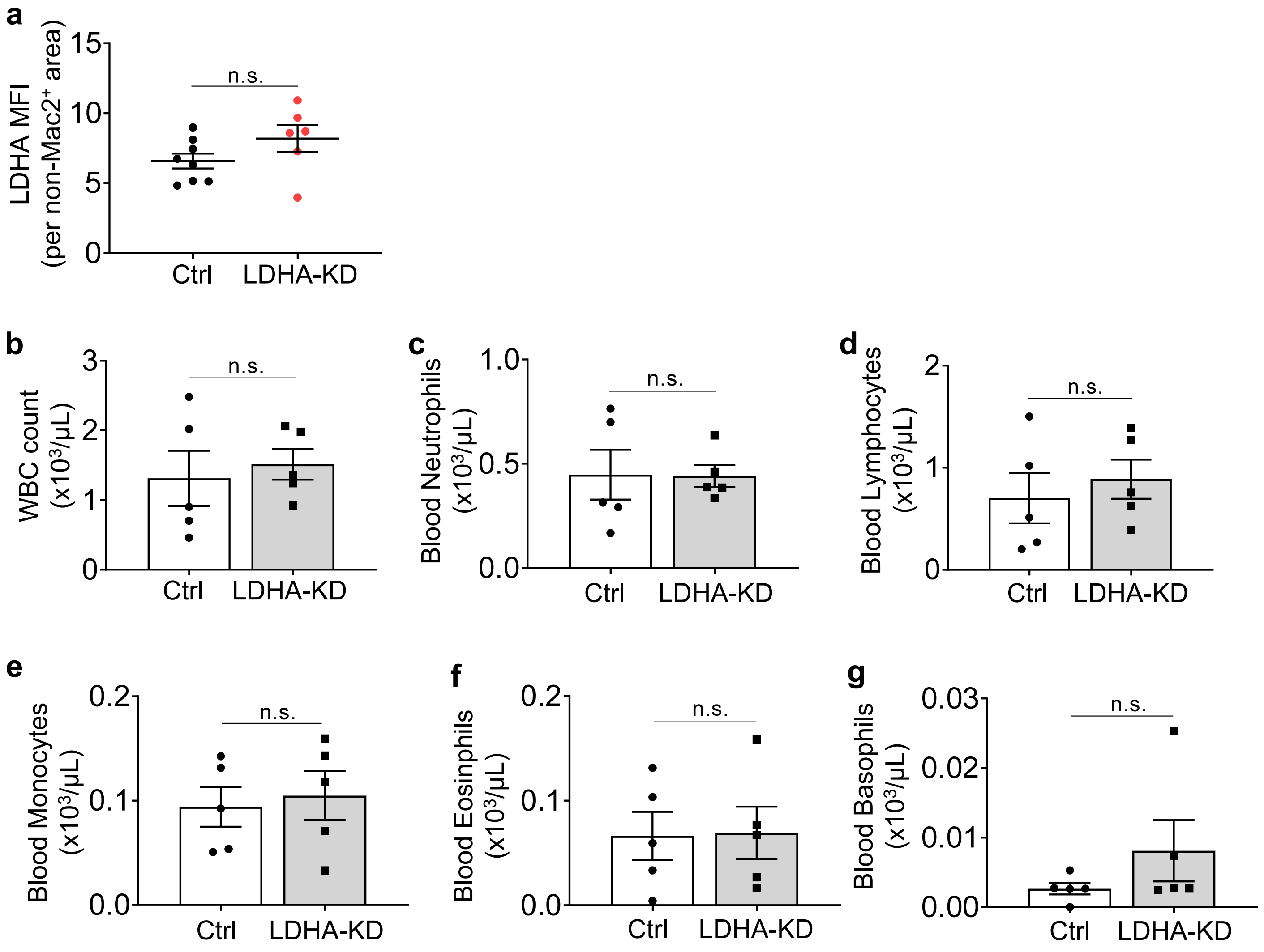


**Extended Data Fig. 2 | Related to Fig. 6: Control and blood counts for the dexamethasone-thymus experiment.**

**a,** Quantification of LDHA MFI in Mac2⁻ areas from Fig. 6b (n=6-8). **b-g,** Counts of blood WBCs, neutrophils, lymphocytes, monocytes, eosinophils, and basophils (n=5). Bars represent means ± SEM. Statistics were performed by student's t-test. n.s., non-significant (*P* > 0.05).

### Supplementary Table

**Supplementary Table 1. Primer sequences used for quantitative RT-PCR analysis**

| Gene | Species | Primer sequence (5' → 3') | |
|---|---|---|---|
| | | Forward | Reverse |
| *Ldha* | Mouse | TGTCTCCAGCAAAGACTACTGT | GACTGTACTTGACAATGTTGGGA |
| *Myc* | Mouse | TGACCTAACTCGAGGAGGAGCTGGAA | AAGTTTGAGGCAGTTAAAATTATGGCT |
| *Gpr132* | Mouse | CTACCTGCGTTTCACCTTTGG | CTGGCAGCTTTGACGAGGAG |
| *Slc16a1* | Mouse | TGTTAGTCGGAGCCTTCATTTC | CACTGGTCGTTGCACTGAATA |
| *Hprt* | Mouse | TCAGTCAACGGGGGACATAAA | GGGGCTGTACTGCTTAACCAG |
| *MYC* | Human | GGCTCCTGGCAAAAGGTCA | CTGCGTAGTTGTGCTGATGT |
| *HPRT* | Human | CCTGGCGTCGTGATTAGTGAT | AGACGTTCAGTCCTGTCCATAA |
